# Cell-density regulated adhesins contribute to early disease development and adhesion in *Ralstonia solanacearum*

**DOI:** 10.1101/2022.11.02.514976

**Authors:** Mariama D. Carter, Devanshi Khokhani, Caitilyn Allen

## Abstract

Adhesins (**adhes**ive prote**ins**) help bacteria stick to and colonize diverse surfaces and often contribute to virulence. The genome of the bacterial wilt pathogen *Ralstonia solanacearum* (*Rs*) encodes dozens of putative adhesins, some of which are upregulated during plant pathogenesis. Little is known about the role of these proteins in bacterial wilt disease. During tomato colonization, three putative *Rs* adhesin genes were upregulated in a Δ*phcA* quorum sensing mutant that cannot respond to high cell densities: *radA* (***R**alstonia* **ad**hesin), *rcpA* (***R**alstonia* **c**ollagen-like **p**rotein), and *rcpB*. Based on this differential gene expression, we hypothesized that adhesins repressed by PhcA contribute to early disease stages when *Rs* experiences a low cell density. During root colonization *Rs* upregulated *rcpA* and *rcpB*, but not *radA*, relative to bacteria in the stem at mid-disease. Root attachment assays and confocal microscopy with Δ*rcpA/B* and Δ*radA* revealed that all three adhesins help *Rs* attach to tomato seedling roots. Biofilm assays on abiotic surfaces found that *Rs* does not require RadA, RcpA, or RcpB for interbacterial attachment (cohesion), but these proteins are essential for anchoring aggregates to a surface (adhesion). However, *Rs* did not require the adhesins for later disease stages *in planta*, including colonization of the root endosphere and stems. Interestingly, all three adhesins were essential for full competitive fitness *in planta*. Together, these infection stage-specific assays identified three proteins that contribute to adhesion and the critical first host-pathogen interaction in bacterial wilt disease.

**Importance:** Every microbe must balance its need to attach to surfaces with the biological imperative to move and spread. The high-impact plant pathogenic bacterium *Ralstonia solanacearum* can stick to biotic and abiotic substrates, presumably using some of the dozens of putative adhesins encoded in its genome. We confirmed the functions and identified the biological roles of several afimbrial adhesins. By assaying the competitive fitness and the success of adhesin mutants in three individual plant compartments, we identified the specific disease stages and host tissues where three previously cryptic adhesins contribute to bacterial success. Combined with tissue-specific regulatory data, this work indicates that *R. solanacearum* deploys distinct adhesins that help it succeed at different stages of plant pathogenesis.

**Research Areas:** Plant Microbiology, Host-Microbial Interactions, Microbial Pathogenesis

## Introduction

Adhesion, or attachment to a surface, transiently or permanently anchors bacteria to a host cell(1–4). After chemotaxis to the host, plant-associated bacteria must attach to the root surface, so adhesion is a critical early step in successful pathogenesis. Further, pathogenic bacteria must adhere to eukaryotic host cells for their type III secretion system to inject the effectors that manipulate host processes to promote bacterial colonization (4). In addition to adhesion, bacteria often participate in cohesion, or interbacterial attachment, which is essential for forming microcolonies and biofilms (5–7). Microcolonies are small collections of a few dozen cells, while biofilms are larger bacterial communities enclosed in a self-produced matrix of extracellular polymeric substances (EPS) that include exopolysaccharides, extracellular DNA, lipids, and proteins (1, 8). Biofilms help bacteria acquire nutrients and provide protection from various environmental stresses, including host defense compounds (9). Thus, pathogenic bacteria need these attachment behaviors to successfully interact with the host and fellow cells and establish disease.

*Ralstonia solanacearum* (*Rs*) causes a lethal vascular wilt disease of diverse plants by infecting hosts through the roots and colonizing their water-transporting xylem vessels. In response to unknown stimuli, *Rs* uses taxis to locate and move to host roots (10). The first point of adhesion is at the root surface, where *Rs* attaches to and forms microcolonies on the surface of root epidermal cells (11, 12). *Rs* enters the root endosphere (interior) through wounds, which can be caused by lateral root emergence, plant parasitic nematodes, or agricultural practices (13). The bacteria then colonize the intercellular spaces of the cortex, where some *Rs* cells form biofilms and reach a high cell density (11, 12, 14). After further invading the root protoxylem through sites where the endodermis is not yet formed or has been breached, *Rs* spreads systemically through the xylem (11, 12). The pathogen attaches to xylem vessel walls, where it forms biofilms and grows to a high cell density. *Rs* populations in stems can reach over 10^10^ cfu/g (1, 15). Wilting results when vessels are occluded by the sheer biomass of the bacterial-EPS matrix and by host defenses like gels and tyloses (16). *Rs* must engage in host and interbacterial attachment at multiple points during this disease cycle, although the specific mechanisms and contributions of individual cell surface proteins to this behavior are not known.

Adhesins (<u>ad</u>hesive prote<u>in</u>s) are surface-exposed or secreted proteins that enable microbial attachment to biotic and abiotic surfaces (4, 5, 17–20). Adhesins are deployed by a wide diversity of symbiotic microbes (21, 22). They are important for the virulence of some plant pathogenic bacteria, contributing to critical interactions with plants, insect vectors, and fellow bacterial cells. For example, a *xadM* adhesin mutant of the rice blight pathogen *Xanthomonas oryzae* pv. *oryzae* is impaired in leaf attachment and biofilm formation *in vitro* and has reduced virulence(23). The xylem-dwelling pathogens *Xylella fastidiosa* and *Erwinia amylovora* require their type I pilus (a multimeric adhesin that forms an appendage on the cell surface) for cell-cell attachment and the initial adhesion required for biofilm formation, respectively (24, 25). *X. fastidiosa* also uses afimbrial adhesins to interact with its insect vectors: *hxfA* and *hxfB* adhesin mutants have reduced attachment to insect foreguts and a lower transmission rate than the wild-type strain (26).

Like many other plant pathogenic bacteria, *Rs* depends on quorum sensing (QS) to regulate genes involved in virulence and other functions in a cell density-dependent manner(27). The global regulator PhcA responds to the Phc QS system and affects expression of over 600 *Rs* genes (15, 28). When the local population of bacteria is relatively small, the QS signal concentration is low and the PhcA regulator is repressed (29, 30). In this low cell density state, *Rs* upregulates dozens of metabolic pathways and nutrient transporters, allowing the pathogen to use a broader pool of nutrient sources when it is away from a host or in the earliest stage of root colonization (15). When *Rs* succeeds in a host and reaches a cell density corresponding to around 5×10^6^ cfu/ml in liquid culture, accumulating QS signal relieves PhcA repression (29, 30). PhcA then represses many catabolic pathways and transporters while simultaneously upregulating virulence factors including exopolysaccharides and plant cell wall-degrading enzymes (15). A *phcA* mutant produces and senses the QS molecule but cannot respond to the QS system. It is thus genetically locked in a low cell density-mimicking state. The *ΔphcA* mutant initially grows faster than wildtype but without virulence factors it quickly fails inside host plants. Profiling the transcriptome of model *Rs* strain GMI1000 during tomato pathogenesis revealed that 14 of the more than 30 putative adhesin genes were differentially expressed in a Δ*phcA* mutant (15). This indicates that much of the *Rs* adhesin repertoire is responsive to cell density and regulated by the QS system via PhcA.

The differential expression of many adhesins in Δ*phcA* suggests that some adhesins play specific roles when *Rs* is at low cell density, presumably corresponding to early stages of disease, while others are needed later in infection when *Rs* is at a higher cell density. We hypothesized that the pathogen deploys and depends on specific adhesins at different points in its life history. But exactly how does *Rs* use these proteins to succeed in plants? We constructed targeted mutants lacking three putative “early” adhesin genes that were upregulated in Δ*phcA*: afimbrial adhesin *radA* (***r**alstonia* **ad**hesin **A**) and two putative collagen-like protein (CLPs) genes *rcpA* (***r**alstonia* **c**ollagen-like **p**rotein **A**) and *rcpB.* Using a suite of *in planta* and *in vitro* assays to quantify bacterial success at specific life stages, we found that these three adhesins contribute to *Rs* attachment, virulence, and fitness in tomato plants. In particular, early adhesins were important for the first critical host-bacterial interaction, attachment to the root surface.

## Results

### *In silico* analysis of the putative adhesins

*rcpA* and *rcpB* are adjacent to each other in the chromosome, possibly in an operon, and are both annotated as collagen-like triple helix repeat-containing proteins and hemagglutinins (Fig 1B). Congruent with this annotation, both proteins were predicted to have CLP (collagen-like protein) middle domains (Fig 1A). RadA is annotated as a YadA-like family protein and surface-exposed adhesin protein. Its name reflects its shared domain architecture with the *Yersinia* trimeric autotransporter adhesin YadA, including multiple YadA-related structural domains like the stalk, head, and anchor (Fig 1A).

**Figure 1.**
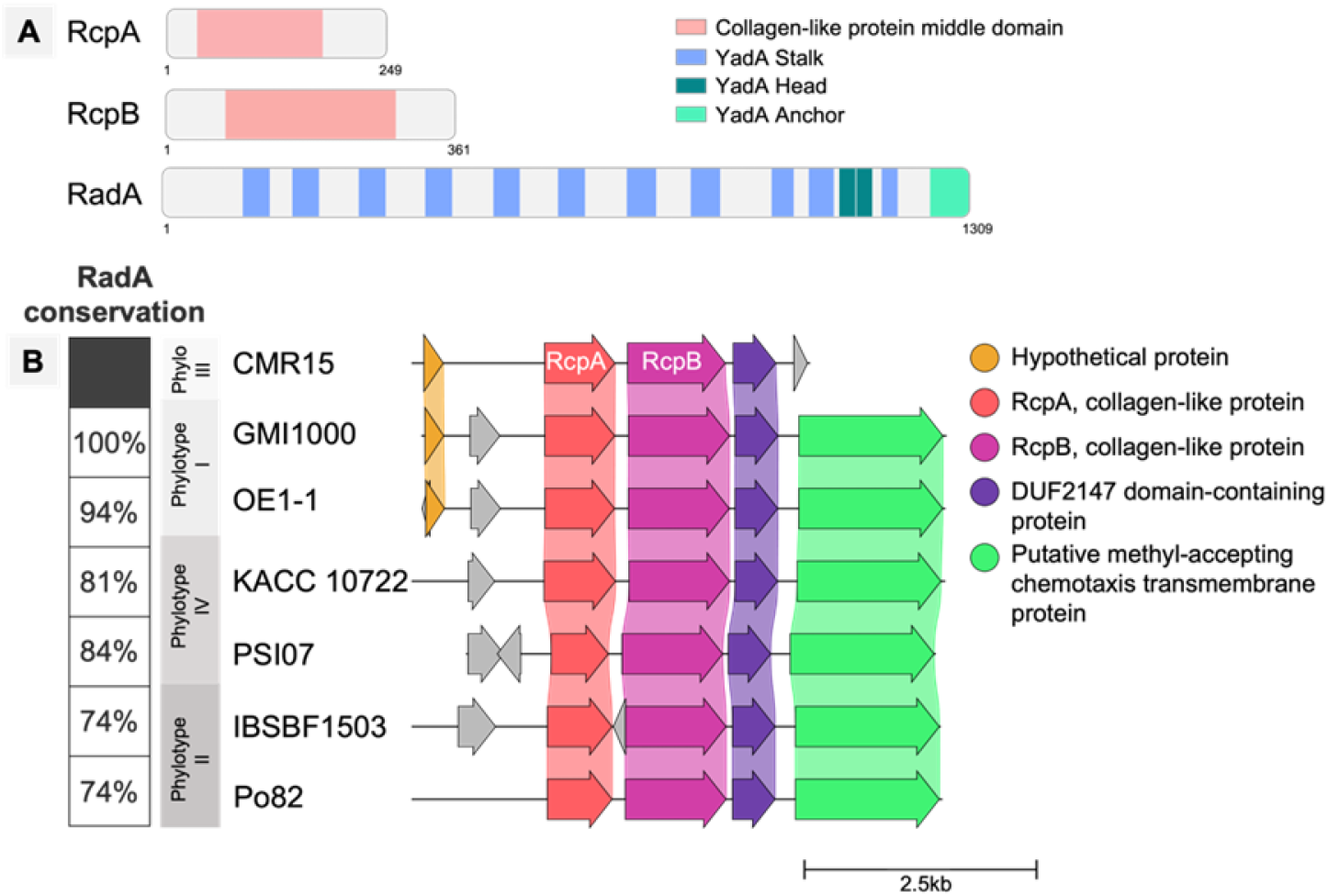
Adhesin protein domain architecture and *rcpA/B* synteny across the RSSC. **A.** The protein families and domains of RcpA, RcpB, and RadA were predicted using the InterPro database. The amino acid sequences were mined for signal peptide sequences with SignalP 6.0. **B.** Clinker was used to generate gene clusters of the RcpA and RcpB genomic region in 7 RSSC strains representing the four phylotypes. Genes marked in the same color have a minimum alignment sequence identity of 0.69.

*Rs* strain GMI1000 is part of the *Ralstonia solanacearum* species complex (RSSC), a large, heterogenous group of strains with a broad geographic distribution and highly variable host ranges. Although phylogenetic analysis divides the RSSC into four phylotypes and three species, all strains colonize plant xylem vessels and cause wilt disease (16, 31, 32). Of 110 RSSC strains analyzed for GMI1000 adhesin homology, 71 have *rcpA*, 54 have *rcpB*, and 54 contain *radA*. Additionally, 47 strains across the four phylotypes encode both *rcpA* and *rcpB*, with a conserved adjacent synteny. Seven of these are shown in Fig 1B.

### RcpA and RcpB gene expression is upregulated during root colonization

Fourteen putative adhesins were differentially expressed when the low-cell-density-mimicking Δ*phcA* mutant grew in the tomato plant (Supplemental Fig S1). Three genes were significantly upregulated relative to in wild-type strain GMI1000: *radA*, *rcpA*, and *rcpB* (2.3, 3.82, and 3.23 log_2_-fold change, respectively) (15). This indicates that the putative adhesins are regulated by the Phc QS system and suggests they may function when the pathogen is at a low cell density, as during soil survival and early root colonization. However, the Δ*phcA* transcriptomic analysis was performed on *Rs* cells inoculated directly into tomato stems where the bacterium experiences very different environmental signals than it does in soil and at the root. Furthermore, because the PhcA regulon is large and complex, we could not rule out pleiotropic effects of the *ΔphcA* mutation.

We therefore used qRT-PCR to determine how these three early adhesins are expressed in a wild-type *Rs* background in other host plant tissues and at varying cell densities. We extracted RNA from *Rs* strain GMI1000 colonizing three distinct compartments of the tomato host: the rhizoplane at low cell density (the site of the first bacterial-host interactions), the root endosphere (the interior of surface-sterilized roots, which contain mixed populations of bacteria at low and high cell densities), and the mid-stem (where *Rs* has grown to high cell densities and formed biofilms).

A gene encoding catabolism of myo-inositol, *iolG*, is required for *Rs* root colonization and is upregulated at low cell density like the early adhesins (15, 33). Expression of *iolG* therefore served as a control to confirm that *Rs* populations sampled from roots were in low cell density regulatory mode. As expected, *iolG* expression was about two-fold (log_2_) higher in *Rs* cells colonizing roots than in *Rs* cells in tomato stems (Fig 2). As a second control we measured expression of *epsB*, a gene involved in biosynthesis of exopolysaccaride (EPS), which is essential for effective stem colonization and is upregulated at high cell density (34). Relative to *Rs* cells from tomato mid-stems, *epsB* expression was quite low in *Rs* cells harvested from root compartments, especially in cells from root surfaces, confirming the QS status of our sampled populations (Fig 2).

**Figure 2.**
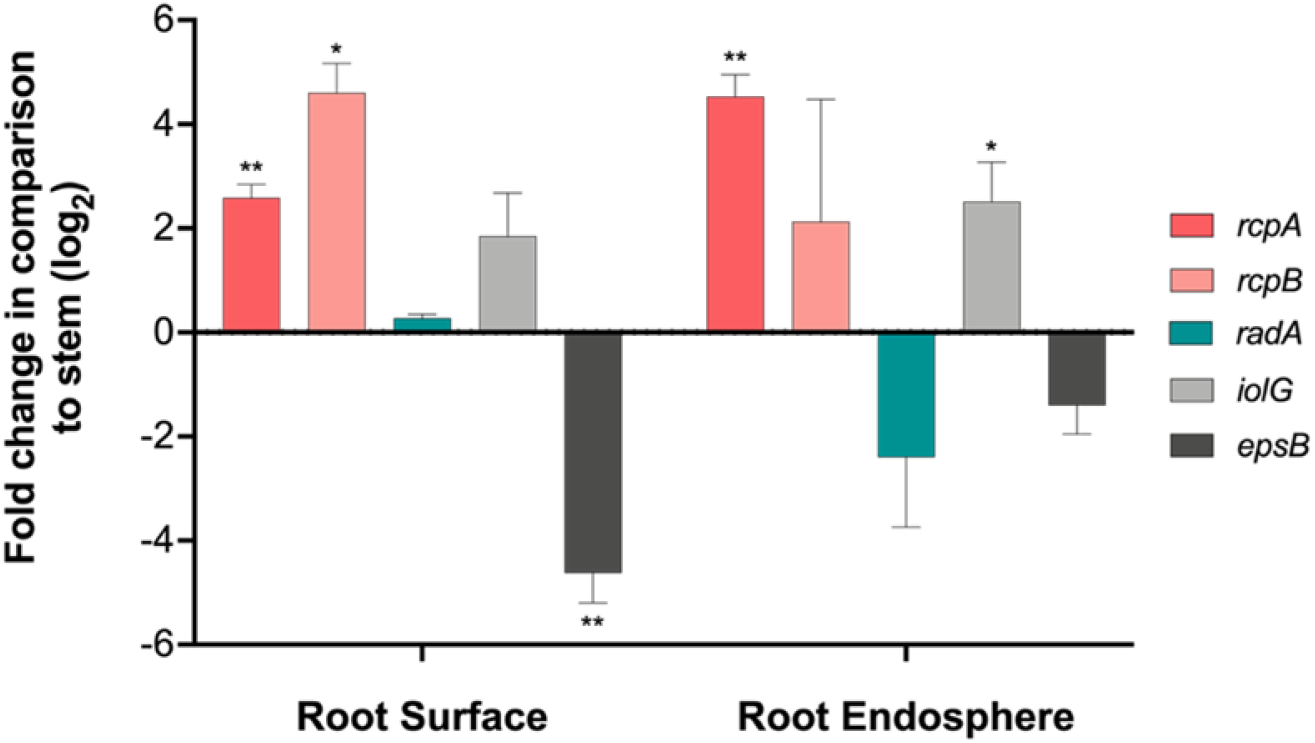
Adhesin gene expression on the root surface and endosphere relative to *Rs* in stems. Five day-old ‘Bonny Best’ tomato seedlings were flood-inoculated with 15 ml (root surface) or 3 ml (root endosphere) of wild-type *Rs* GMI1000 at 5×10^6^ cfu/ml suspended in chemotaxis buffer. To measure gene expression of *Rs* cells colonizing the rhizoplane (root surface), roots were harvested 6 hpi, gently washed in water once, and cut from the hypocotyl. Total RNA was extracted from pools of 40 to 50 homogenized roots. To measure gene expression in *Rs* cells colonizing the root endosphere, RNA was extracted from pools of surface-sterilized and homogenized roots harvested 48 hpi. Data shown are from two biological replicates, each containing 6 technical replicates. To measure gene expression in *Rs* colonizing tomato mid-stem, RNA was extracted from mid-stems of 24-day old plants 3 days after petiole inoculation with 2 μl of 1×10^6^ cfu/ml. Data shown reflect three biological replicates consisting of 4 pooled plants. The targeted gene expression on the root rhizoplane and in the root endosphere is compared to the stem. The fold change is presented on a

Both *rcpA* and *rcpB* were expressed at higher levels in *Rs* cells harvested from tomato root surfaces than in bacteria growing in the stem (Fig 2). In bacteria from the root endosphere, *rcpA* was significantly upregulated and *rcpB* expression trended upwards, suggesting these proteins function in root colonization. Under these conditions *radA* expression did not differ across the three host compartments, although *radA* expression trended downwards in the root endosphere (Fig 2). Thus, qRT-PCR analysis confirmed that *Rs* upregulates the putative adhesin genes *rcpA* and *rcpB* during root colonization relative to later in wilt pathogenesis when *Rs* has reached high cell densities in the stem. The putative adhesin gene *radA* was expressed at comparable levels in all three plant compartments at low and high cell densities.

### *RadA*, *RcpA* and *RcpB* are required for competitive fitness *in planta*

To further explore functions of these adhesins, we constructed targeted *Rs* deletion mutants *ΔradA* and *ΔrcpA/B*. Because their genomic proximity and similar expression levels suggest an operon and they encode proteins that share motifs, *rcpA* and *rcpB* were deleted together for functional characterization. We used a holistic virulence assay to determine if loss of these early adhesins affected wilt disease progress. When unwounded tomato plants were soil-soak inoculated with the adhesin mutants, *ΔrcpA/B* caused wilt disease indistinguishable from that of the wild-type parent strain while *ΔradA* had a minor but significant virulence defect early in disease (Fig 3A and 3B).

**Figure 3.**
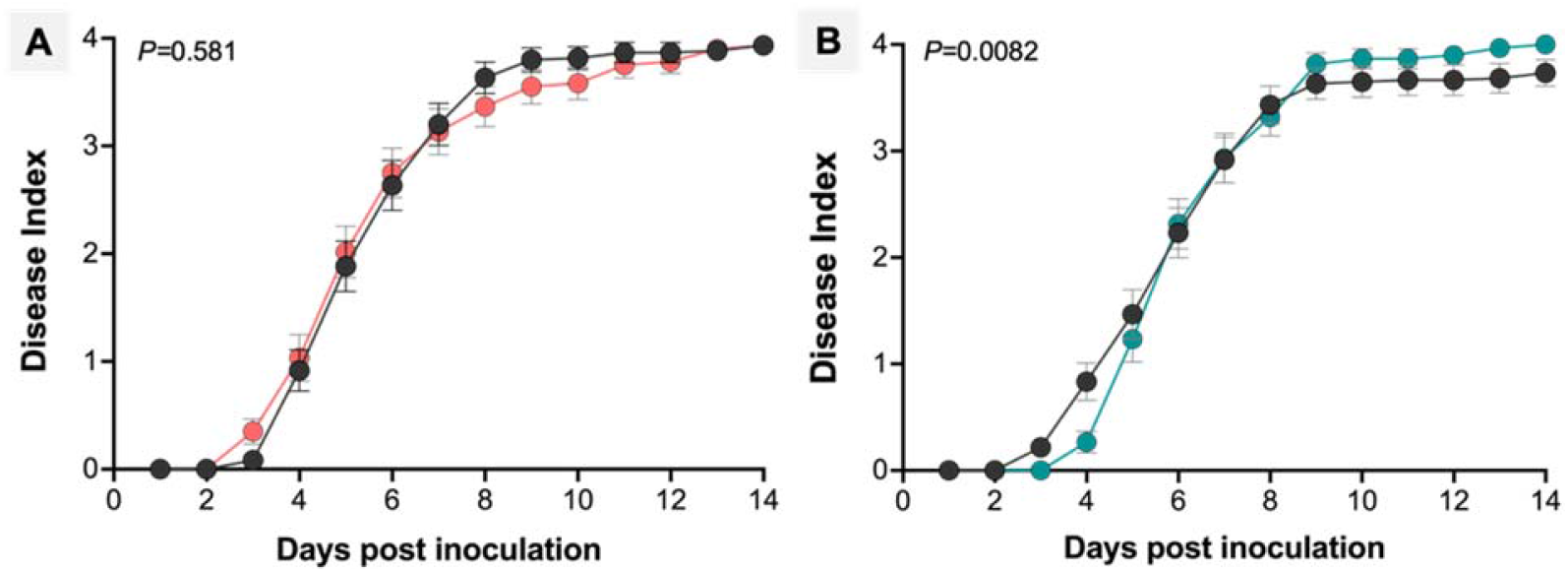
The Δ*radA* mutant has a virulence defect following soil-soak inoculation, while Δ*rcpAB* does not. Unwounded three-week-old tomato plants were drenched with 1×10^8^ cfu/g soil of wild-type strain GMI1000 (black symbols), *ΔrcpA/B* (red symbols) (**A**), or *ΔradA* (teal symbols) (**B**). Plants were rated daily for wilt symptoms over 14 days using a disease index scale ranging from 0-4 (0: no leaves wilted, 4: 75-100% of leaves wilted). Data shown reflects three biological replicates, each with 20 tomato plants (A, *P*=0.581; B, *P*=0.0082, Repeated Measures ANOVA).

Because the soil-soak disease assay measures only the final output of wilt symptom development, we used more focused experiments to determine mutant fitness and behavior in specific host environments. To assess the contribution of RadA, RcpA, and RcpB to aboveground plant colonization, we quantified *Rs* populations in tomato mid-stems harvested 4 or 5 days after soil-soak inoculation. The Δ*rcpA/B* double mutant reached a smaller mean population size in the stem than the wild type after soil soak inoculation, but this trend was not significant (Fig 4A). When bacteria were directly inoculated into tomato stems via a cut petiole, the Δ*rcpA/B* and Δ*radA* mutant colonized plants as well as the wild-type parent (Supplemental Fig S2B, Supplemental Figure S3B).

**Figure 4.**
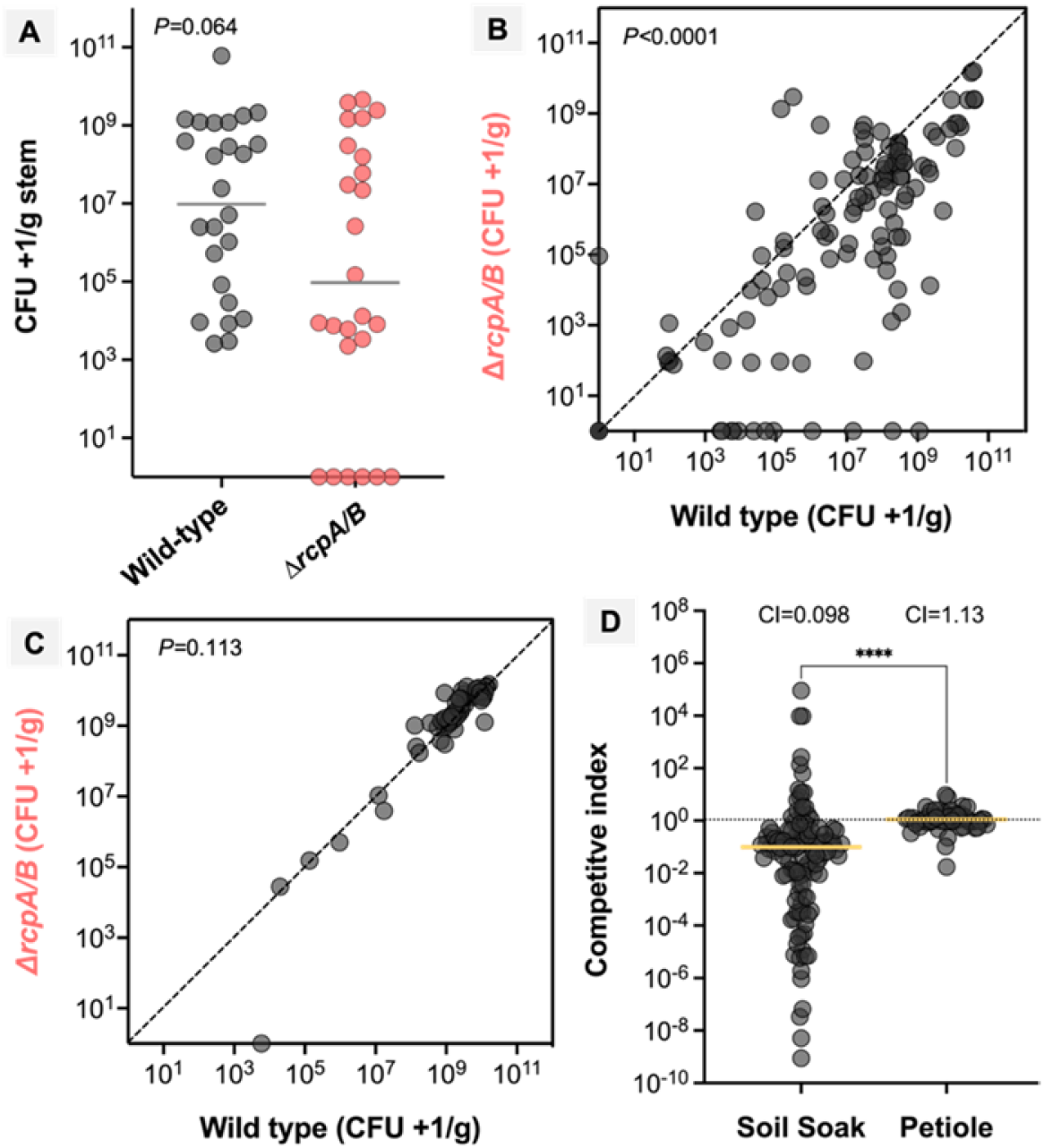
The Δ*rcpA/B* mutant *in planta* competitive fitness defect was rescued by direct stem inoculation. 21-day old tomato plants were individually inoculated (**A**) or co-inoculated by soaking soil (**B**) with 1:1 suspensions of wild-type GMI1000-Km and ΔrcpA/B-Gm or via cut-petiole inoculation (**C**), and the population size of each strain was determined by dilution plating ground mid stem sections on appropriate selective media. The individual strain soil soak experiments were harvested 4 dpi. Data shown are two biological replicates with 10-15 plants (Mann-Whitney test, *P*=0.064). The grey bar is the geometric mean. For soil soak co-inoculation experiments, tomato mid-stems were sampled 4 to 7 dpi. The wild-type strain outcompeted the mutant at all time points (Wilcoxon test, *P*<0.0001). The experiment was done twice with 20 technical replicates for the co-infection treatments. At 3 days after co-inoculation via cut petiole, mutant and wild-type populations were not different (Wilcoxon test, *P*=0.113). The experiment was done twice with 30 technical replicates for the co-infection treatments and 15 technical replicates for the individual inoculations. (**D)** For both the petiole and soil soak experiments, the competitive index (CI) was calculated as the ratio of mutant cfu/g to wild-type cfu/g stem. A CI=1, shown as dotted black lines, means the wild-type and Δ*rcpA/B* populations are equal in the stem (Mann-Whitney test, *P*<0.0001). Yellow bars

Co-inoculating a mutant with the wild-type strain can reveal subtle but biologically important defects in pathogen fitness. We therefore measured competitive fitness of the adhesin mutants during tomato colonization by soil-soak inoculating plants with a 1:1 ratio of antibiotic marked wild-type and mutant strains. The population size of each strain was measured in mid-stems harvested 4 to 7 dpi. Loss of the CLPs RcpA and RcpB together significantly decreased *Rs* competitive fitness (Fig 4B, 4D). The wild type outcompeted Δ*rcpA/B*, consistently reaching larger population sizes across all time points (Supplemental Fig S2A). To determine if this fitness defect was the effect of cumulative disadvantage over the disease process or was specific to the stem environment, antibiotic marked Δ*rcpA/B* and wild-type cells were directly co-inoculated into stems via a cut petiole to bypass the steps of root entry, root colonization, and early xylem colonization. Interestingly, the *rcpA/B* mutant retained full competitive fitness under this condition (Fig 4C and 4D, Suppl Fig S2B). Additionally, Δ*rcpA/B* had no fitness defect when competitively colonizing the seedling root endosphere for 24 h (Suppl Fig S2C). These results suggest that in addition to aiding attachment to seedling roots, Δ*rcpA/B* may function in early attachment to the tomato xylem walls.

*Rs* cells lacking afimbrial adhesin RadA were less affected in competitive fitness. After soil soak co-inoculation with wild-type, the population of Δ*radA* was similar to that of wild-type 4 days after inoculation and significantly lower on days 5, 6, and 7 (Supplemental Fig S3A). Combining data from days 4 to 7 revealed Δ*radA* had a slight but significant competitive defect (Fig 5B and D). Unlike Δ*rcpA/B* however, co-inoculating wild type and Δ*radA* into a cut leaf petiole did not rescue the mutant’s competitive fitness (Fig 5C and 5D, Supplemental S3B). This defect was specific to the stem environment because Δ*radA* had no fitness defect in competitive colonization of the root endosphere (Supplemental Fig S3C). Together, these experiments showed that RadA contributes to pathogen behaviors in the stem, which could explain its virulence defect. While RcpA and RcpB were not required for bacterial wilt virulence under tested conditions, deleting the early adhesins did reduce *Rs* competitive fitness *in planta*.

**Figure 5.**
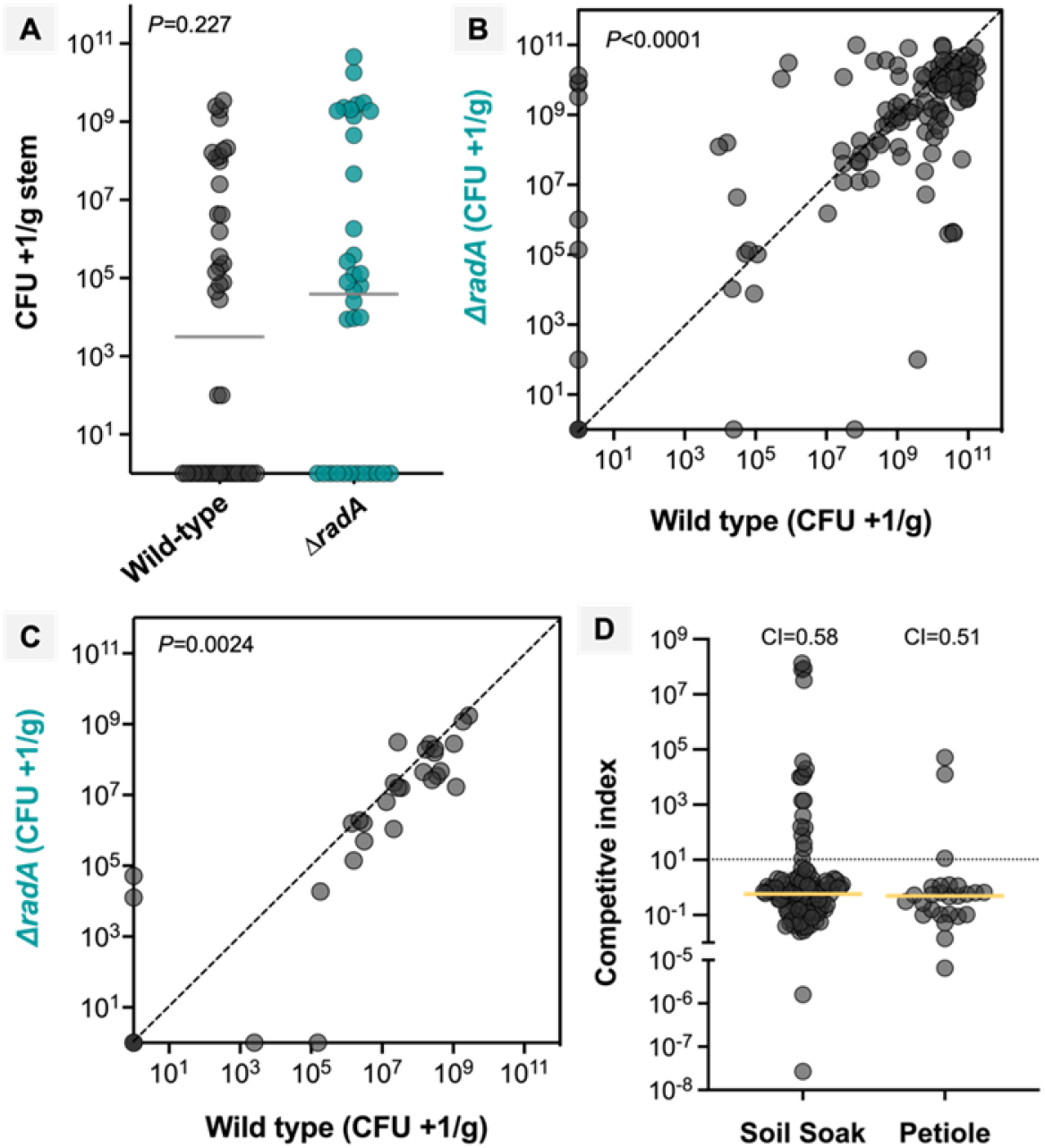
The Δ*radA* mutant has reduced competitive fitness in the stem. 21-day old tomato plants were individually inoculated (**A**) or co-inoculated by soaking soil (**B**) with 1:1 suspensions of wild-type GMI1000-Km and ΔradA-Gm or via cut petiole inoculation (**C**), and the population size of each strain was determined by dilution plating ground mid stem sections on appropriate selective media. The individually soil soaked experiments were harvested 4-5 dpi. The data shown are from three biological replicates with 10 to 15 plants (Mann-Whitney test, *P*=0.227). The grey bar is the geometric mean. For soil soak co-inoculation experiments, tomato mid-stems were sampled 4 to 7 dpi. The wild-type strain outcompeted the mutant on days 5 to 7 (Wilcoxon test, *P*<0.0001). The experiment was done twice with 20 technical replicates for co-infection treatments. Petiole inoculation did not rescue Δ*radA* competitive fitness defect (Wilcoxon test, *P*=0.0024). The experiment was done twice with 24 technical replicates for the co-infection treatments and 10 technical replicates for the individual inoculations. (**D).** For both the petiole and soil soak experiments, the competitive index (CI) was calculated as the ratio of Δ*radA* cfu/g to wild-type cfu/g stem. The dotted black line marks a CI=1, meaning the wild-type and Δ*radA* populations are equal in the stem. Yellow bars indicate the median. The median CI is below one for both the petiole and soak soil experiments, which are similar (Mann-

### The early adhesins help *Rs* attach to abiotic surfaces and form biofilm

Despite the intense shear force generated by vascular flow, *Rs* bacteria anchor themselves to the xylem wall and form biofilms, which require both adhesion to a surface and cell-cell attachment (1, 15). To assess the role of the early adhesins in this complex attachment behavior, we visualized GFP-expressing strains forming biofilms on glass with confocal microscopy. After 2 days of static culture on glass well slides, the supernatant was removed with gentle aspiration and the bottom of the chamber was imaged (Fig 6A-6C). To remove cells that were not tightly adhered to the glass, the wells were gently washed once with water and imaged again (Fig 6D-6F). Before rinsing, the wild-type cells formed a typical bacterial biofilm consisting of an irregularly dense layers with three-dimensional structure. In contrast, Δ*radA* and Δ*rcpA*/B formed smooth layers of cells on the slide with a flat surface topography. Surprisingly, a single gentle wash removed most of the mutant strain cells, indicating that they were not tightly attached to the glass. While washing also removed some wild-type bacteria, many thick aggregates remained attached to the slide. These findings indicate that the early adhesins contribute to a normal biofilm architecture and specifically mediate adhesion to abiotic surfaces.

**Figure 6.**
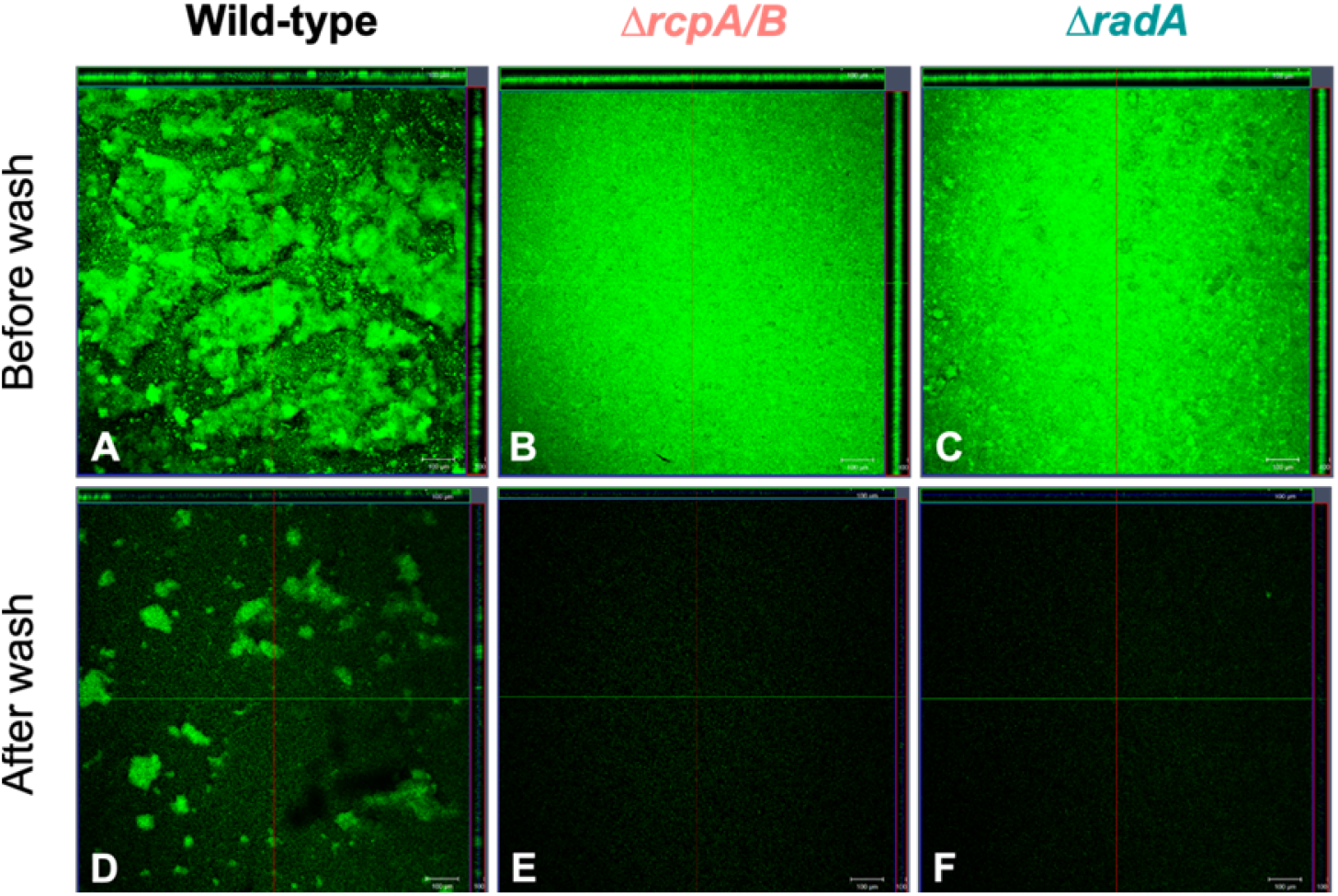
RadA and RcpA/B contribute to adhesion to abiotic surfaces. 500 μl cultures of GFP-expressing wild-type and mutant *Rs* were grown in glass chamber slides. Two dpi, the supernatant was removed and the biofilm that formed on the bottom of the chamber was imaged using confocal microscopy (**A-C**). To remove cells that were not specifically adhered to the glass, the wells were gently washed once with 500 μl of sterile water and imaged again (**D-F)**. A representative orthogonal view of the biofilm is shown. The experiment was done twice with 4 technical replicates per biological

### RadA, RcpA, and RcpB contribute to root attachment

Before it can reach the stem *Rs* must colonize the root interior, or endosphere. Microscopic analysis found that *Rs* colonized the outer and inner root cortex by two days after inoculation (11). To measure the contribution of the early adhesins to infection of the root cortex, we inoculated tomato seedlings with *Rs* and after 48 h, surface sterilized the roots and quantified the endosphere bacterial population. Wild-type *Rs* and the adhesin mutants all reached a similar average population size of 10^7^ cfu/g root (Fig 7A). Similar results were observed at 24 hpi (Supplemental Fig S2C). This suggests that RadA, RcpA, and RcpB do not aid in colonization of the root interior, a life stage when many *Rs* cells have reached high cell densities (33).

**Figure 7.**
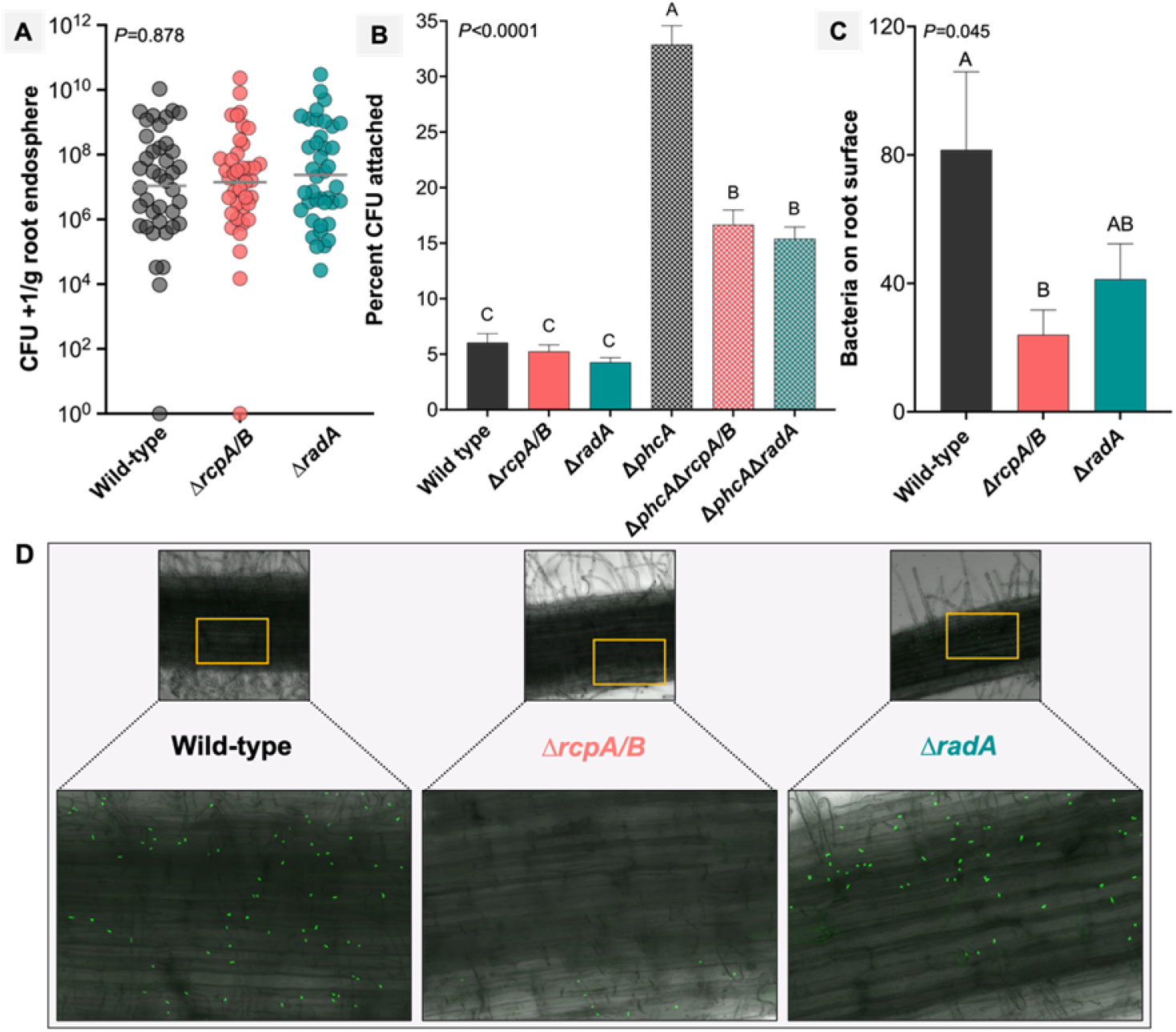
RadA and RcpA/B help *R. solanacearum* adhere to roots, but do not contribute to root endosphere colonization. **A.** Four-day old tomato seedling roots were inoculated with 10,000 cells (10 μl of 1×10^6^ Cfu/ml) of *Rs* wildtype, Δ*phcA*, or adhesin mutant strains. Roots were incubated for 48 h after which they were surface sterilized, homogenized, and bacteria colonizing the root interior were quantified by serial dilution plating. Each dot represents four pooled roots; horizontal gray bar indicates the geometric mean. Experiments were repeated three times with 10 to 20 technical replicates (Kruskal-Wallis test, *P*=0.878). **B.** To access attachment to the root surface, roots were harvested after 4 h, gently washed to remove unattached cells and processed. Bacterial populations recovered from roots were standardized to the initial inoculum amount. Data shown represent three independent biological replicate experiments, each containing 5 to 10 technical replicates per treatment. Technical replicates consisted of four pooled roots. Different letters indicate significance groups (*P*<0.0001, ANOVA). **C** and **D.** Seedling roots were inoculated with GFP-expressing *Rs* strains. After 4 hours, roots were washed once, placed on cover glass, and visualized with a confocal microscope. The number of fluorescent bacteria on the root surface were counted manually at three locations on each root and measured on four roots per strain (*P*=0.045, ANOVA). Representative images of bacterial attachment near the root hypocotyl are shown.

At the very beginning of disease *Rs* adheres to the root surface, where it forms microcolonies before exploiting wounds for root entry. We hypothesized that this initial attachment step is facilitated by *Rs* adhesins that are expressed under low cell density conditions, such as the bacterium experiences in soil. Wild-type *Rs* cells attach to host roots at a low frequency, with only about 3% of inoculated cells strongly attached to root epidermal cells after 2 h (35). These small numbers make it difficult to detect minor attachment defects. We therefore quantified root attachment of *ΔradA* and *ΔrcpA/B* mutants in both the wild-type *Rs* background and in the *phcA* QS mutant. The *ΔphcA* mutant hyper-attaches to tomato root surfaces and it is thus a useful positive control.

Bacteria at a low cell density were pipetted onto the surface of axenic tomato seedling roots. After 4 h, roots were gently washed to remove unattached bacteria, then ground and serially dilution plated to quantify the root-associated bacterial population. This assay revealed that early adhesins contributed to root attachment (Fig 7B). In the wild-type background, neither mutant displayed a significant attachment defect, although both consistently trended towards lower root attachment. As expected, the Δ*phcA* mutant attached more than 5 times as well as the wild type. Deleting either *radA* or *rcpA/B* from this hyper-attaching mutant background reduced *Rs* attachment by nearly 50% (Fig 7B).

To further assess the role of the early adhesins in root attachment, we directly visualized GFP-expressing bacteria on tomato roots using confocal microscopy. This independent method also found that deleting early adhesins reduced *Rs* root attachment, with a slightly different result (Fig 7C and 7D). The number of root-attached *ΔradA* mutant cells trended lower than that of wild-type, while the *ΔrcpA/B* double mutant was significantly reduced in root attachment and there was no difference between the *ΔradA* and *ΔrcpA/B* mutants. Together, these experiments showed that Rs adheres to host roots in part because of RadA and because of RcpA, RcpB or both.

## Discussion

From *Marinomonas primoryensis* living in multispecies Antarctic biofilms to *Xanthomonas* living on rice and citrus leaves, bacteria interact with their environment physically using adhesins (36–39). Adhesins are often key virulence factors that help pathogenic bacteria form biofilms that protect them from host defense compounds and anchor them in flowing environments like xylem vessels (40, 41). The genome of xylem-inhabiting *Ralstonia* strain GMI1000 encodes dozens of putative adhesins, but their function in pathogenesis is largely unknown.

This study focused on three of these adhesins: *radA*, *rcpA*, and *rcpB*. RadA was so named because it contains several domains present in YadA, an autotransporter adhesin common in *Yersinia* spp., including a coiled stalk, head domain, and anchor (42, 43). YadA, a virulence factor in *Y. enterocolitica*, mediates interactions between the bacterium and host cell surface glycans, and promotes autoaggregation (cell to cell attachment) (43–45). Its similarity to YadA suggests that RadA is an adhesin of the same class and might be secreted by the type V secretion system (19). RcpA and RcpB are much smaller proteins, at 249 and 361 AA, respectively, and both contain a central collagen-like protein (CLP) middle domain. Like mammalian collagen, bacterial CLPs have a characteristic Gly-X-Y repeat that generates a triple helical structure (46). Although plants do not contain collagen, CLPs mediate interactions between cells of the plant growth-promoting rhizobacterium *Bacillus amyloliquefaciens* (47, 48). RcpA and RcpB are annotated as having CLP middle domains and contain multiple characteristic GXY repeats, but these repeats are not in the contiguous clusters of three or more typically associated with bacterial CLP proteins. This suggests that RcpA and RcpB are unlikely to form the stable triple helices that are a core feature of animal collagen and typical bacterial CLPs. Further, the predicted structures and sequence of RcpA and RcpB offer no clues about their cellular localization. Additional experiments are needed to determine if these three adhesins are present on the cell surface and directly engage in adhesion.

Genes for RadA, RcpA, and RcpB are present in dozens of RSSC strains across the four phylotypes, representing vast geographic and host diversity. While *radA* is present in over 50 genomes, it is notably missing from phylotype III (African) strains (Fig 1B). The synteny of the *rcpA* and *rcpB* genomic region is present in multiple strains as well. The fact that many RSSC strains make these attachment proteins is consistent with their biological importance and suggests they play similar roles in strains across the complex and in diverse plant hosts.

The *Rs* PhcA quorum sensing regulatory system represses all three adhesin genes under high cell density conditions; consistent with that, a Δ*phcA* mutant overexpresses them in tomato mid-stems. Based on these observations, we initially hypothesized that the pathogen depends on RadA, RcpA, and RcpB very early in disease because in nature *Rs* populations probably reach quorum only in the nutrient-rich and confined habitats inside plants. Analysis of wild-type *Rs* gene expression in multiple host environments demonstrated that *rcpA* and *rcpB* were upregulated during early root colonization, as predicted. However, *radA* expression levels were similar whether *Rs* was on root surfaces, inside the root cortex, or colonizing stems, suggesting that the contribution of *radA* spans the disease cycle. This result also demonstrates that signals other than cell density regulate *radA* during pathogenesis. This is not surprising because bacteria integrate many environmental and plant cues into complex regulatory networks.

We measured adhesin mutant phenotypes using a diverse set of assays that each had advantages and limitations. The holistic soil soak inoculation disease progress assay mimics the natural *Rs* infection process by forcing the bacterium to first chemotax through soil to host roots, then attach to and colonize root surfaces, invade the xylem and finally multiply there to cause the wilt symptoms that are the assay output. To ensure that the wild-type strain reliably kills all or most plants, soil-soaking employs an atypically high *Rs* soil population of 10^8^ cfu/g soil. This overwhelming inoculum level can compensate for quantitative defects early in the infection process, and thus obscure the impacts of subtle but biologically significant virulence factors.

For example, an *Rs iolG* mutant, which can’t catabolize the myo-inositol that is abundant in root exudates, is as virulent as wild-type following soil-soak inoculation (33). However, like *ΔrcpA/B*, *ΔiolG* is significantly delayed in the early stages of root colonization (33). Similarly, in the xylem pathogen *Xanthomonas fuscans* subsp. *fuscans* the nonfimbrial adhesin YapH is essential for initial adhesion to biotic and abiotic surfaces even though a *yapH* mutant was as virulent as the wild type in a bean leaf dip assay (49). In contrast, *ΔradA* had a small but significant virulence defect in soil-soak assays, consistent with a function beyond root adhesion and also with the fact that *radA* is expressed at similar levels on root surfaces, root endospheres, and in mid-stem xylem vessels.

We evaluated the contributions of the adhesins more narrowly by measuring bacterial population sizes at intermediate points in disease development, in root endospheres and in the midstem. When inoculated individually, both adhesin mutants colonized root endospheres and tomato mid-stems as well as the wild-type strain, although midstem population sizes of Δ*rcpA/B* did trend lower than those of the wild-type strain. All strains reached similar population sizes if they were directly inoculated into mid-stems, notably lacking the defect trend observed for Δ*rcpA/B* following soil-soak inoculation.

Co-inoculations can detect subtle reductions in bacterial fitness and isolate the points in the disease cycle where a trait is relevant. When we forced wild-type and adhesin mutant strains to compete for limiting resources like host attachment sites, wild-type *Rs* consistently outcompeted Δ*rcpA/B*. These adhesins likely do not contribute to the later stages of stem colonization because introducing a mixture of these two strains directly into xylem rescued the colonization defect of Δ*rcpA/B*. Although we cannot rule out the possibility that the rapid growth and large populations following petiole inoculation could conceal a subtle defect, this result suggests that RcpA and RcpB are required for full competitive fitness during disease development at points somewhere between colonization of the root interior and late colonization of the mid-stem. *Rs* cells likely cycle between a low and high cell density mode as the pathogen spreads through the host. Thus, RcpA and RcpB may contribute to early xylem attachment when *Rs* cells first invade a vascular element. Unlike Δ*rcpA/B*, *ΔradA* was not restored to full fitness by petiole inoculation, suggesting that RadA may help with xylem adhesion at mid-stage disease. The fact that co-inoculation with the wild-type strain cannot rescue the fitness defects suggests the mutants are defective in adhesion to the host rather than in cohesion between bacterial cells. Such defects are challenging to measure directly, especially in the xylem, but might be revealed with microscopy of differentially labeled competing strains or by using eventual plant mutants that lack adhesin binding sites.

*Rs* cells form biofilms in xylem and possibly in root intercellular spaces. Diverse *Rs* mutants that do not form normal biofilms *in planta* have virulence defects, including those lacking EPS, extracellular DNAses, and the PhcA regulator (1, 14, 15). Both adhesin mutants failed to develop normal wild type complex biofilm architecture on glass slides. While this biofilm formation is on an abiotic surface, it does suggest that both mutants can autoaggregate but are impaired in adhesion to surfaces. Thus, RadA and RcpA and RcpB together contribute to the initial attachment step of biofilm formation, consistent with roles in attachment to both root surfaces and possibly xylem walls. It would be informative to compare the structures formed in tomato xylem by wild type *Rs* to those formed by the two adhesin mutants.

In all these disease stages, *Rs* is likely at a high cell density or in heterogeneous populations of mixed cell densities across host microenvironments. Both in planktonic liquid culture and in cells recovered from infected tomato stems, *Rs* is at a high cell density, meaning that PhcA-dependent virulence factors are upregulated when the population is >10^7^ cells/ml or cells/g stem tissue (13, 34, 50). *Rs* quickly reaches and exceeds that cell density *in planta* and did so in our plant assays. Plant pathogenic bacterial biofilms can also rapidly reach a high cell density mode in restricted environments where the quorum sensing molecule can accumulate. Aggregates containing as few as 13 *Pseudomonas syringae* cells on a dry leaf surface reached a state of quorum sensing induction(51). Therefore, the negative results from our root colonization and stem colonization assays reinforce that RcpA and RcpB function early in disease at the root surface and possibly at the xylem surface when *Rs* is at a low cell density.

We found that the three afimbrial adhesins contribute to root adhesion, the critical first step in bacterial wilt pathogenesis. Deleting either *radA* or *rcpA/B* from the *ΔphcA* mutant reduced its hyper-attachment to tomato roots. Additionally, confocal microscopy revealed that *Rs* mutants lacking both RcpA and RcpB suffered a defect in attachment to tomato roots, while Δ*radA* trended towards lower attachment but was not significantly different from wild type. Neither adhesin deletion completely abolished attachment, suggesting that additional factors contribute to adhesion and to the notably sticky phenotype of *ΔphcA*. A triple mutant lacking all three genes and a quadruple mutant in Δ*phcA* unfortunately gave variable results, so we were unable to assess the effects of this additional deletion.

Like many plant-associated bacteria, *Rs* uses the type IV pilus, a complex multi-proteinaceous appendage, to attach to roots (52). The *Rs* type IVa pilus is critical for tomato infection and for polar attachment to roots(53). Similarly, the type IVb pilus is required for virulence, particularly in the early stages of disease, and for *in vitro* biofilm formation (54, 55). Treating *Rs* strain UW55I with mannose and fucose reduced bacterial attachment to tomato roots, providing indirect evidence that sugar-binding proteins (lectins) could mediate host root-pathogen interactions (56). Extracellular polysaccharide, a non-proteinaceous compound secreted by the pathogen, was also important for adherence to the root surface (35). To our knowledge, this is one of the first demonstrations that non-pilus and non-fimbrial adhesins contribute to root-bacterium interactions in bacterial wilt disease. Congruent with their expression on the rhizoplane, *rcpA* and *rcpB* function in the earliest stage of disease when the bacteria are at a low cell density. Although it was not uniquely upregulated during root colonization, *radA* contributed to root attachment to the same degree as *rcpA* and *rcpB* together. Thus, *Rs* appears to deploy a diversity of proteinaceous and non-proteinaceous cell surface structures to mediate this critical step in host colonization.

## Methods

### Bacterial strains and culture conditions

A list of bacterial strains used in this research can be found in Supplemental Table S1. *Rs* strains were grown from −80°C glycerol stocks or water stocks on rich medium containing casamino acids, peptone, and glucose (CPG) and tetrazolium chloride (TZC), supplemented with antibiotics when needed (15 mg L^−1^ gentamicin and 25 mg L^−1^ kanamycin), and grown at 28°C for 48hrs (57). Overnight cultures of *Rs* were grown in CPG broth or Boucher’s Minimal Media supplemented with 0.2% glucose (w/v) (BMM, pH 7.0) in a 28°C shaking incubator(58). *Escherichia coli* (*E. coli*) strains were grown at 37°C on Luria-Bertani (LB) media, supplemented with kanamycin and gentamicin when necessary.

### Mutagenesis

Cloning, restriction digestion, sequencing, and PCR were performed using standard methods(59, 60). DNA sequencing was performed at the University of Wisconsin-Madison Biotechnology Center and oligonucleotide primers were synthesized by Integrated DNA Technologies. Gibson assembly (New England BioLabs, Ipswich, MA) was used to create in-frame deletion constructs with the primers listed in Supplementary Table S1. Deletion constructs were introduced into the chromosome of wild-type *R. solanacearum* GMI1000 and Δ*phcA* by double homologous recombination via electroporation or natural transformation followed by *sacB* counter-selection to create mutant strains GMI1000Δ*rcpA/B*, GMI1000Δ*radA*, Δ*phcA*Δ*rcpA/B*, and Δ*phcA*Δ*radA* (61). For strain competition assays, we marked the mutants in the wild type background with gentamycin by integrating pRCG-GWY into the chromosome *attTn7* site. To generate GFP-expressing adhesin mutants, GMI1000-gfp genomic DNA was used in a natural transformation of Δ*rcpA/B* and Δ*radA* to produce Δ*rcpA/B*-gfp and Δ*radA*-gfp.

### *In vitro* biofilm formation

The biofilm assay was performed as described by Tran (2016). Briefly, overnight cultures of GFP-expressing *Rs* strains were grown in CPG broth, pelleted, washed, and resuspended in fresh CPG media. The cultures were standardized to 10^8^ cfu/ml, diluted into to 500 μl of 10^7^ cfu/ml in the wells of an 8-well Nunc Lab-Tek II chamber cover glass (Thermo Fisher Scientific). The cover glass was covered with the provided cap and incubated at 28°C for 72 h. The spent media was replaced with fresh CPG every day. After the three-day incubation period, the media was removed, and the biofilms imaged with a Zeiss LSM 710 laser scanning confocal microscope at the University of Wisconsin-Madison Newcomb Imaging Center. The wells were then washed once with 500 μl of sterile water and imaged a second time. Each assay contained 3-4 technical replicates and the was repeated two times.

### Seedling root assays

Tomato seeds (bacterial wilt-susceptible cv. ‘Bonny Best’) were surface sterilized with 10% bleach for 10 min and 70% ethanol for 5 min then grown on 1% water agar with a Whatman Grade 1 85 mm filter paper disk (Cytiva, Marlborough, MA) for four days. *Rs* inoculum was prepared by growing overnight in CPG at 28C with shaking. Cultures were pelleted, washed with water, diluted to 1×10^6^ cfu/ml, and allowed to adjust to a low cell density regulatory mode for 2 h in BMM. This suspension was pelleted and 10ul suspension of the bacteria resuspended in water to a final concentration of 1×10^6^ cfu/ml was pipetted directly onto the surface of four-day-old seedling roots. After 4 h, 2-cm root sections were excised, gently washed in water, and blotted dry. Four roots were pooled per sample, ground, serially diluted, and plated to quantify cfu attached per cm of root. The proportion of cells that attached to the root surface was calculated by normalizing to the population size of the initial inoculum, as determined by dilution plating (62).

For confocal microscopy, plates of four-day-old roots were flooded with a *Rs* suspensions at low cell density. After incubating for 4h, roots were gently washed in water, blotted dry, and placed on a glass slide and covered with cover glass for visualization. The roots were imaged with a Leica SP2 AOBS laser scanning confocal microscope at the Ohio State University and a Zeiss LSM 710 laser scanning confocal microscope at the University of Wisconsin-Madison Newcomb Imaging Center. A 488nm laser was used to excite *Rs* produced GFP and emitted fluorescence collected between 500 and 519nm. A 405 nm laser was used to capture root autofluorescence and emitted fluorescence collected between 430 and 470 nm. Cfu on the root surface were counted from maximal projections images.

For root endosphere colonization, four-day old seedling roots were inoculated with 10,000 cells from each unmarked *Rs* strain (10 μl of 1×10^6^ cfu/ml). Roots were incubated for 48 h at room temperature under a 12 h light cycle. Following colonization, the roots were surface sterilized for 30sec with 10% bleach, 30 sec with 70% ethanol, and then washed four times in sterile water. The roots are cut from the hypocotyl, pooled 4 roots per sample, weighed, ground, and serially diluted. The data reflects three biological replicates, each with 10-20 technical replicates. For competitive root colonization, roots were inoculated with antibiotic marked strains, wild-type GMI1000-Km, ΔradA-Gm or ΔrcpA/B-Gm individually or in a 1:1 ratio of the wild-type and mutant strain. Roots were harvested 24hpi, processed, and serially dilution plated on CPG plates amended with kanamycin or gentamycin to distinguish the mutant and wild-type populations.

## Tomato stem assays

Wilt-susceptible ‘Bonny Best’ tomatoes were inoculated by drenching the soil with a bacterial suspension to a concentration of 1×10^8^ cfu/g soil, as previously described (62). Briefly, 21-day old plants were inoculated with the *Rs* strains. Plants were grown in a growth chamber with a 12 h light cycle at 28°C and rated daily for wilt symptoms over 14 days. A 0-4 disease index scale was used: 0, no leaves wilted; 1, 1–25% leaves wilted; 2, 26–50% leaves wilted; 3, 51–75% leaves wilted; and 4, 76-100% leaves wilted. The experiment was repeated three times for each strain with 20 plants per replicate.

To access stem colonization, three-week old tomato plants were soil soak inoculated with antibiotic marked strains, wild-type GMI1000-Km, Δ*radA*-Gm or Δ*rcpA/B*-Gm. Plants were also co-inoculated with a 1:1 mixture of the wild-type and adhesin mutant strains at a concentration of 1×10^8^ cfu/g. Four to 7 days post-inoculation, 100 mg mid-stem sections were excised, homogenized in 900 μl of sterile water using a PowerLyzer (MO BIO, Carlsbad, CA) and serially dilution plated on CPG plates amended with kanamycin or gentamycin. Experiments were repeated twice with 20 plants per replicate for the co-inoculations and 10-15 plants per replicate for the individual inoculations.

For petiole inoculation experiments, the petiole of the oldest true leaf was cut and inoculated with antibiotic marked strains individually or co-inoculated. A 2ul droplet of bacterial suspensions at 1×10^6^cfu/ml was placed onto the cut surface as previously described (62). Bacterial population sizes were measured by serial dilution plating as described above onto CPG plates with kanamycin or gentamycin to quantify the two strains. Experiments were repeated twice with 30 plants per replicate for the co-inoculations and 15 plants per replicate for the individual inoculations.

### Statistical Analyses

Graphing and statistical analyses were done using GraphPad Prism 8.0. The specific statistical tests used for experiments are provided in figure legends.

## Acknowledgements

This material is based upon work supported by the National Science Foundation Graduate Research Fellowship under Grant No. 1650114. The authors gratefully acknowledge Sarah Swanson, University of Wisconsin-Madison Department of Botany, and Jonathon M. Jacobs, Ohio State University Department of Plant Pathology, for their technical help with confocal microscopy. We also thank Corri D. Hamilton, University of British Colombia Department of Microbiology and Immunology, for designing the *iolG* qPCR primers.

